# Total variation of quantitative phase map reveals cellular Young’s modulus

**DOI:** 10.1101/2025.09.08.674874

**Authors:** Zhenghua Wang, Zebin Wang, Deyu Sun, Yin Li, Yujie Nie, Jialin Shi, Renjie Zhou, Lianqing Liu, Min Long, Yongliang Yang

**Affiliations:** The State Key Laboratory of Robotics and Intelligent Systems, Shenyang Institute of Automation, Chinese Academy of Sciences, Shenyang 110016, China; The University of Chinese Academy of Sciences, Beijing 100049, China; School of Electrical and Control Engineering, Shenyang Jianzhu University, Shenyang, 110168, China; Shenzhen BayJayRay Biomedical Technology Co., LTD., Shenzhen 518000, China; Department of Biomedical Engineering, The Chinese University of Hong Kong, Shatin, New Territories, Hong Kong SAR, China; Department of Endocrinology, Southwest Hospital Affiliated to Army Medical University; Chongqing, 400000, China

## Abstract

Collective cell migration is a fundamental physiological process involved in wound healing, development, and tissue regeneration. Though the role of the environmental stiffness in collective cell migration has been extensively investigated, the effects of cellular stiffness have been less studied due to the lack of high-throughput in situ methods for characterizing cellular stiffness. Here, we characterize the cellular Young’s modulus in situ with a large field of view and in real time using use quantitative phase microscopy. We found that standard deviation and total variation of the phase are inversely related to the cellular Young’s modulus, while the total variation of phase has finer mechanical resolution. Integrating the total variation of phase with cell segmentation algorithms, we efficiently analyzed the cellular Young’s modulus for cells in a monolayer. Using this system, we found the cells at the wound frontier are much softer than that of the cells in the inner region of monolayer in *in vitro* wound healing assay. At single cellular level, the Young’s modulus of leader cells, boundary cells, and inner cells are significantly distinct from each other. To conclude, our method would help elucidate the essential role of cellular stiffness in collective cell migration and has the potential to further benefit the progress of mechanobiology.

## Introduction

Collective cell migration plays an essential role in a wide range of biological processes, such as embryogenesis, tissue regeneration, and cancer metastasis ^1,2^. In epithelial collective cell migration, cells in an integrated monolayer migrate together while maintaining biophysical and biochemical interactions. During migration, cell behaviors are influenced both by their intrinsic stiffness and the environmental stiffness ^3^. For example, gradients of substrate stiffness guide the collective cell monolayer migration ^4,5^. However, compared to the effects of environmental stiffness on collective cell migration, the role of cellular stiffness has been less studied due to the limitations of instruments. It requires to measure cellular stiffness in a large area in situ manner with fast tempera resolution, which has not been realized yet.

Various methods for measuring cellular stiffness have been developed. The most reliable and widely used method is force indentation of atomic force microscopy (AFM). In this method, an AFM cantilever indents the cell being measured and the cellular stiffness is calculated from cantilever deformation data using the Hertz model ^6,7^. This technique characterizes cellular stiffness in situ with nano-newton and nanometer resolution ^8^. To acquire an image containing both spatial and mechanical properties of a cell, the AFM tip sequentially scans the cellular area, which process lasts tens of minutes or even longer (Figure 1a). Thus, it is not suitable for high-throughput measurements with high temporal resolution. Magnetic tweezers, optical tweezers, and optical elastography can deform cells and measure cellular stiffness in a high throughput manner ^9–11^. All these methods, however, require placing cells in solution and deforming them significantly, rather than in situ measurements, such as migrating on a substrate. The recently developed brillouin microscopy can probe cellular mechanical properties in situ with high spatial resolution without damaging the cells ^12,13^. The required high pump powers and long acquisition time (increasing the signal-to-noise ratio of the measurement) in Brillouin microscopy limits its implementation of imaging highly dynamic cellular behaviors ^13,14^. Therefore, to investigate the role of cellular stiffness in collective cell migration, a new instrument that can measure cellular stiffness in high-throughput in situ manner with single cell resolution is still needed.

**Figure 1.**
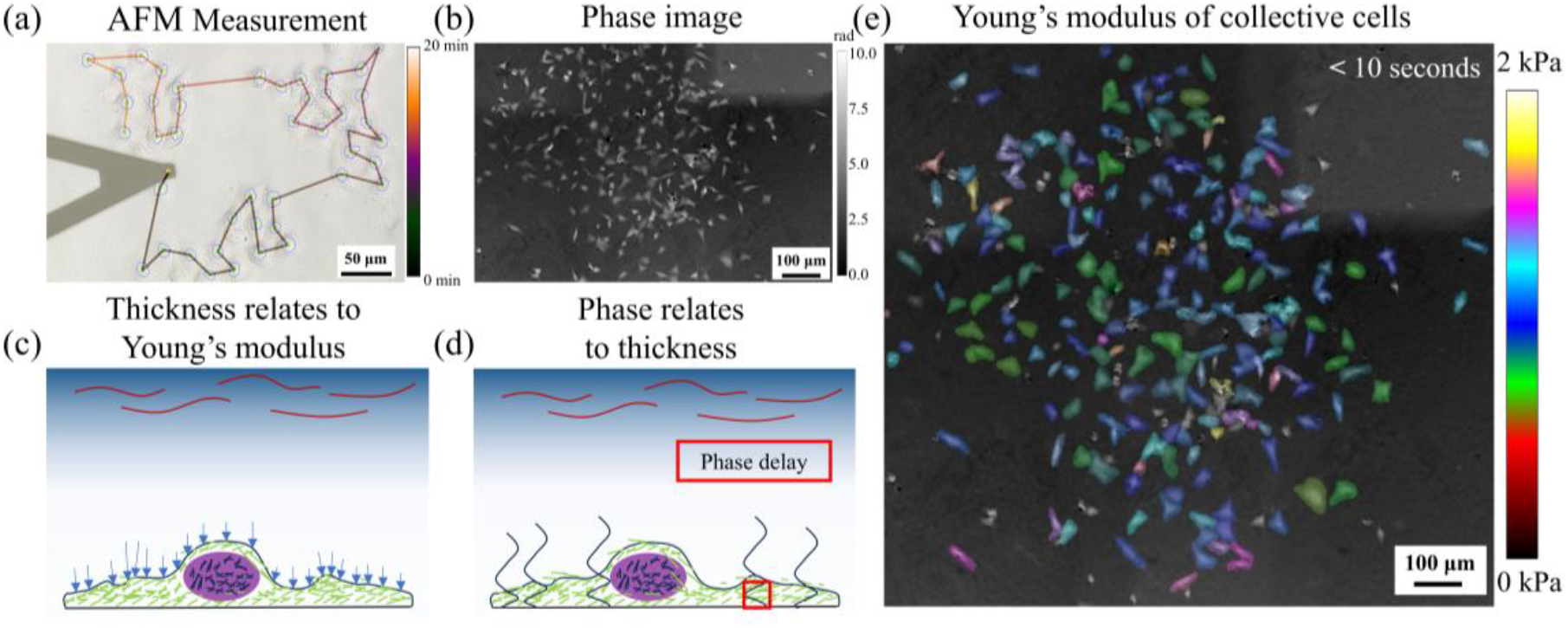
>Scheme of using phase for high-throughput collective cellular Young’s modulus measurement. (a) Time-consuming AFM measuring process. (b) High-throughput phase image. (c) Thickness link to cellular Young’s modulus. (d) Phase link to cellular Young’s modulus. (e) Real-time Young’s modulus characterization of collective cells.

Interferometric quantitative phase microscopy (iQPM) offers new opportunities for high-throughput measurements of cellular mechanical properties in situ (Figure 1b) ^15^. iQPM splits its optical beam into a reference beam and a sample beam. Due to the difference of refractive index between the sample and the medium, the sample beam experiences an optical phase delay. Quantitative measurement this phase delay has been widely used to image the cellular shear stress and organelle dynamics in vitro, and thermal deformations in vivo ^16–18^. It has also been adopted for cancer detection based on the mechanical properties of cancer cells ^19–21^. Prior research demonstrated that membrane fluctuations measured by iQPM can provide insights into the mechanical properties of erythrocytes (red blood cells) ^22^. By imaging the spatial deformation of phase delay, iQPM has the potential to characterize cellular mechanical properties without deforming cells by using the disorder strength of phase ^23,24^. Disorder strength calculated from phase indicated the length of fiber-like cytoskeletal structures, which is correlated with cellular stiffness ^25^. This characterization of disorder strength makes iQPM promising for high-throughput measurements of cellular mechanical properties ^26^.

The mechanical properties of cells are mainly determined by the amount and structure of various cytoskeletons, such as intermediate filament and F-actin ^27,28^. According to cellular tensegrity, the cytoskeleton senses mechanical forces and plays a major role in cell stiffness ^29–31^. Considering the molecules of the medium are constantly colliding with the cell membrane with a statistically constant force, we hypothesized that the cellular stiffness is linked to fluctuations in cellular thickness, which accompanied with changes in cellular volumes (Figure 1c) ^32^. Here, we introduced two metrics to represent the fluctuations: standard deviation and total variation. We first observed an inverse relationship between cellular thickness fluctuations and Young’s modulus in AFM images of height and Young’s modulus. When light passing through the cell body, the cellular thickness, together with cellular refractive index, determines the optical phase delay (Figure 1d) (details are in MATERIALS AND METHODS Standard Deviation and Total Variation). By assuming the cellular refractive index as a constant in low resolution, the cellular Young’s modulus is inversely related to phase fluctuation and can be characterized a a large area to achieve high throughput. The phase fluctuations, especially the total variation of phase, might represent the distribution of cytoskeleton, like disorder strength of phase ^25^. We treated cells with Y-27632 (a ROCK inhibitor ^33,34^) to manipulated the cytoskeleton, and the phase fluctuations characterized the corresponding cellular Young’s modulus changes. To further characterization in a high-throughput manner, we used low-magnificent objective lens to enlarge field of view, and the total variation of phase is powerful for high-throughput characterization.

We combined the total variation of phase and cell segment method (ASF-YOLO ^35^) to rapidly characterize the cellular Young’s modulus (within 10 seconds) as shown in Figure 1e and Figure S1. After observing collective cell migration using the total variation of phase, we found that the spatial distribution of cellular Young’s modulus in monolayer is inversely relate to the distance from frontier, and trend to a constant. The gradient vectors of cellular Young’s modulus near edge exhibits significant directionality. Notably, the gradient vectors of Young’s modulus are oriented toward the leader cells. We further used total variation of phase to analyze the collective cells movements for 6 hours and observed the change of Young’s modulus of a leader cell, a follow cell, and an inner cell. These results demonstrated that the developed method would help elucidate the essential role of cellular stiffness in collective cell migration and has the potential to further benefit the progress of mechanobiology.

## RESULTS

### Cellular Young’s modulus inversely relates to fluctuations of cellular thickness

We scanned the local area of a C2C12 cell using AFM to establish the relationship between the variation of thickness and cellular Young’s modulus (Figure 2a). Areas with more fiber-like structures were stiffer (Figure 2b). As presented in Figure 2c and 2d, standard deviation of thickness and total variation of thickness were inversely related to cellular Young’s modulus. The soft cellular area exhibited large fluctuations, while the hard cellular area showed small fluctuations. Presenting these two metrics in relative manners showed a similar trend as shown in Figure 2e and Figure 2f respectively. Using AFM, we further scanned the whole cell as presented in Figure 2g and Figure 2h. Standard deviation of thickness (Figure 2i) and total variation of thickness (Figure 2j) in images were also inversely related to Young’s modules (Figure 2h) of the whole cells. Because the areas of nucleus have less cytoskeleton structures, they are softer than the cytosol full of cytoskeleton. We observed a similar trend in the two metrices as percentages, as shown in Figure 2k and Figure 2l. To further investigate the relationship between Young’s modulus of a cell and its cytoskeleton, we stained F-actin and nuclei of the same cells as shown in Figure 2m. In the cytoskeleton aggregated area, we observed similar fiber-like stiff areas (high Young’s modulus area in Figure 2h), corresponding to high Young’s modulus values. And according to those stiffer areas, we observed lower thickness fluctuations (Figure 2i-l).

**Figure 2.**
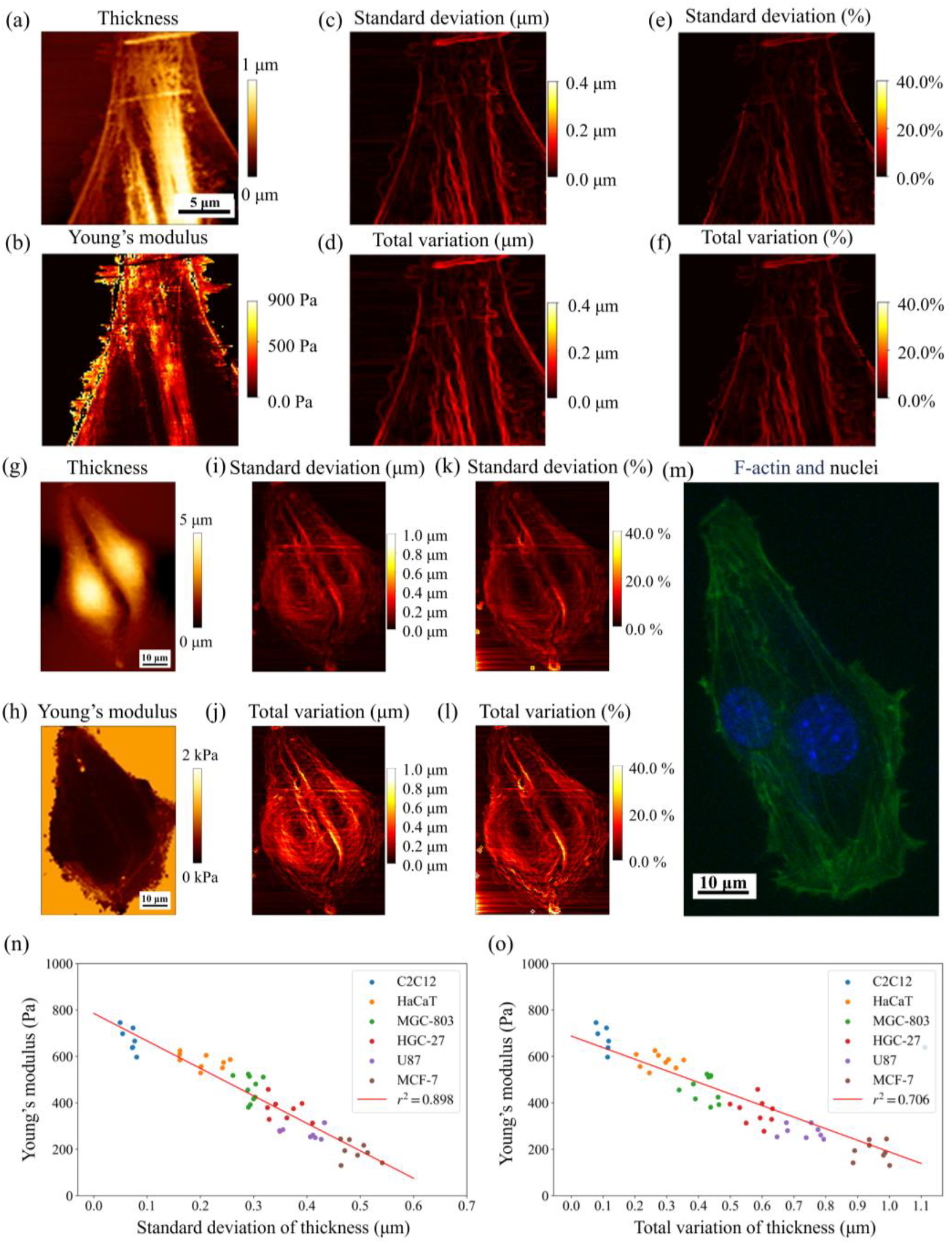
>Standard deviation and total variation of thickness are inversely related with the Young’s modules of cells. (a) The thickness in a local area of a cell. (b) The cellular Young’s modulus in local area. (c) The standard deviation of thickness in local area. (d) The total variation of thickness in local area. (e) The percentages of standard deviation of thickness in local area. (f) The percentages of total variation of thickness in local area. (g) The thickness in the whole cell. (h) The Young’s modulus in the whole cell. (i) The standard deviation of thickness in the whole cell. (j) The total variation of thickness in the whole cell. (k) The percentages of standard deviation of thickness in the whole cell. (l) The percentages of total variation of thickness in the whole cell. (m) The fluorescence image of F-actin fiber in green and nuclei in blue. (n) The inverse relationship between Young’s modulus and standard deviation of thickness. (o) The inverse relationship between Young’s modulus and total variation of thickness.

To quantify this relationship, we scanned six types of cells using AFM (C2C12, HaCaT, MGC-803, HGC-27, U87, and MCF-7). More AFM images were shown in Figure S3. Considering the substrate effects inherent in AFM measurements, we limited our analysis to measurements where the Young’s modulus were ≤1500 Pa. We calculated the average Young’s modulus and two metrics of thickness for each cell and plotted the results as scatter points in Figure 2n and Figure 2o. While both metrices exhibited an inverse relationship, total variation of thickness displayed a smaller absolute value of the slope.

### Standard deviation and total variation of phase also inversely relate to cellular Young’s modulus

Scanning cells using an AFM cantilever is low-throughput and time-consuming. Because the optical phase delay is mainly influenced by the cellular thickness in low resolution high-throughput imaging (details are in **MATERIALS AND METHODS** Standard Deviation and Total Variation), we considered the feasibility of using optical phase delay to indicate the variations of cellular thickness. As shown in Figure 3a and Figure 3b, the phase values in nuclei areas were larger than that of other areas, which is similar as the thickness. We hypothesized that fluctuations of phase were also inversely related to the Young’s modulus of cells. We further calculated the total variation of thickness and phase, as shown in Figure 3c and Figure 3d. Total variation of thickness had similar pattern to that of total variation of phase. In the cellular areas with large thickness fluctuation, the phase fluctuation values were also large. We further observed more types of cells using iQPM and calculated their total variation of phase and standard deviation of phase (Figure S4). The total variations of phase were larger than the standard deviation of phase, consistent with Figure 2 in AFM results. This was due to that standard deviation calculated the fluctuation of a region comparing to the mean value of that region.

**Figure 3.**
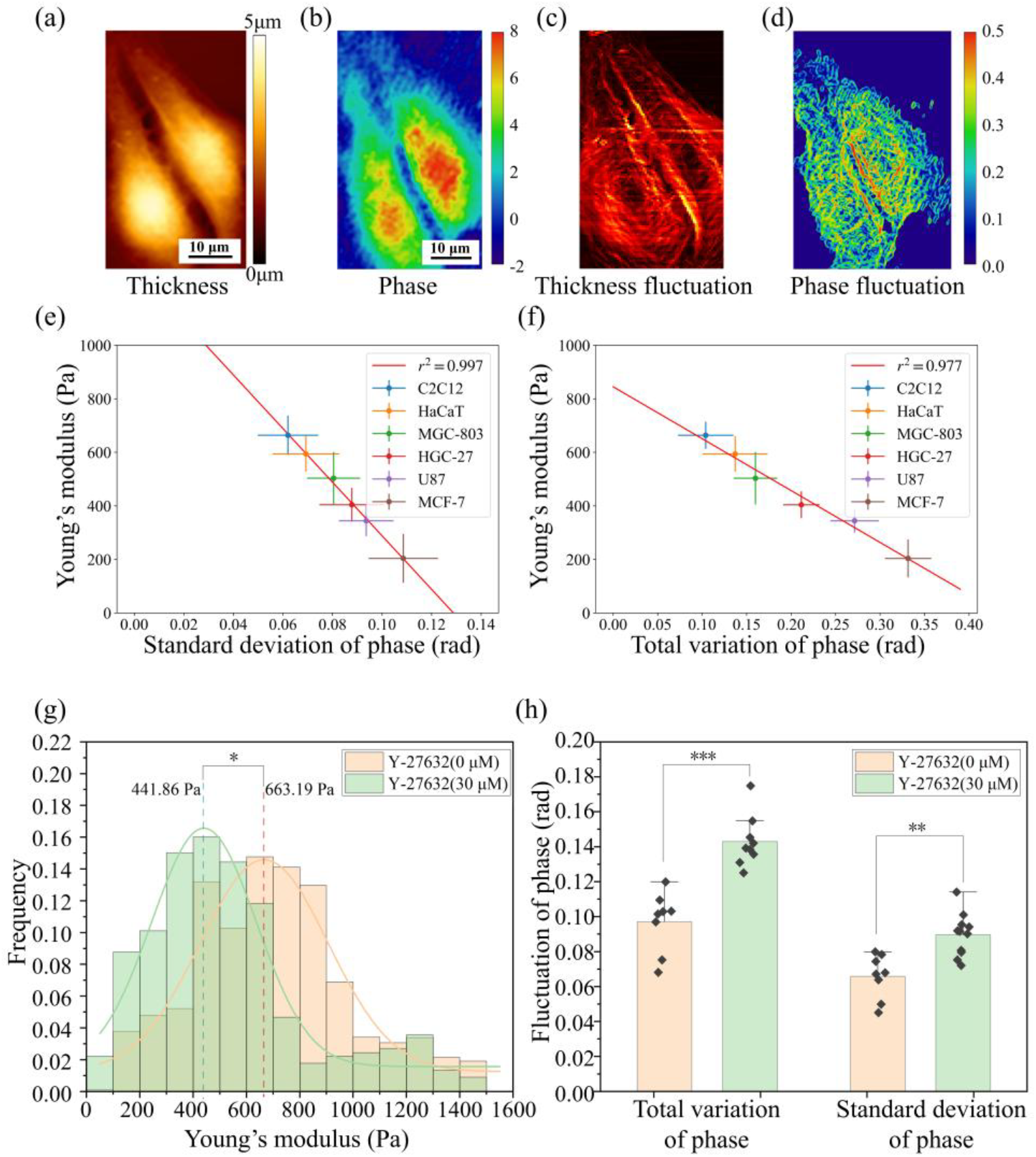
>Standard deviation of phase and total variation of phase are inversely related with the Young’s modules. (a) The thickness of C2C12 cells. (b) The phase of C2C12 cells. (c) The total variation of thickness of C2C12 cells. (d) The total variation of phase of C2C12 cells. (e) The inverse relationship between Young’s modulus and standard deviation of phase. (f) The inverse relationship between Young’s modulus and total variation of phase. (g) The reduction of Young’s modulus caused by Y-27632. (h) The reflecting increasement of both two metrices of phase according to the reduction of Young’s modulus.

To demonstrate the relationship between the Young’s modulus of cells and phase fluctuations, we averaged the metric values of about 10 cells for each type. The relationship between Young’s modulus and both two fluctuation metrices of phase were linearly fitted, as shown in Figure 3e and Figure 3f. The two metrics proposed here were inversely related to cellular Young’s modulus. Interestingly, total variation of phase displayed a smaller absolute value of the slope (Figure 3f), indicating better resolution of representing cellular Young’s modulus using total variation of phase. If we exclude HaCaT, the total variation of phase could classify the other five cell types, which is not the case in the standard deviation. Overall, these results demonstrated the inverse relationship between Young’s modulus and the two metrics of phase.

To further explore the relationship between fluctuations of phase and the intracellular cytoskeleton structures, we manipulated the cytoskeleton structures using Y-27632, a ROCK inhibitor ^33,34^. Demonstrated by the AFM results in Figure 3g, the C2C12 cells softened after Y27632 treatment. We then subjected C2C12 cells to identical conditions and imaged them using the iQPM and calculated the two metrices of phase. Shown in Figure 3h, the two proposed metrics of phase in the Y-27632-treated cells increased compared to those in the control group, which is consistent with our hypothesis. Furthermore, the difference in total variation of phase is more statistically significant than that of standard deviation of phase.

### The proposed total variation of phase performs well in low-resolution phase images for high-throughput applications

To characterize the cellular Young’s modulus in a high throughput manner during collective cell migration, the iQPM should have a large field of view, which reduces its spatial resolution. Thus, we used an objective lens (OL) with low magnification for a large field-of-view (FOV) and low resolution (Figure 4a). The clearly visible subcellular structures using 40× OL were gradually invisible in images using 10x OL. Meanwhile, the physical distance between neighboring pixels was larger than that in high-resolution phase images. Thus, the cellular thickness represented by phase values between neighboring pixels changed significantly, which increases the fluctuations of phase. The correlation between cellular Young’s modulus and the two proposed metrics of phase in 10× OL was shown in Figure 4b and c respectively. The standard deviations of phase of some cell types overlapped with each other, whereas the total variations of phase of these cell types were distinguishable. More iQPM images and inverse relationship of both two metrices in 20× OL were shown in Figure S5 and Figure S6. To further increase the FOV to about 1cm^2^, we used the 10× OL for scanning the collective cells with overlap to observe large FOV phase image. After 9 times (3×3 scanning) imaging, the large FOV phase image was shown in Figure 1b. We used ASF-YOLO to segment each cell in the phase image and calculated the mean total variation of phase for each cell ^35^. By integrating the mean total variation of phase of these cells with the inverse relationship between cellular Young’s modulus and total variation of phase, we characterized the Young’s modulus of many cells as shown in Figure 1e. For well visualization, the characterization of cellular Young’s modulus of collective cells was combined with the phase image and amplitude image to yield a clear result (Figure S1). The time cost including scanning and computing is less than 10 seconds.

**Figure 4.**
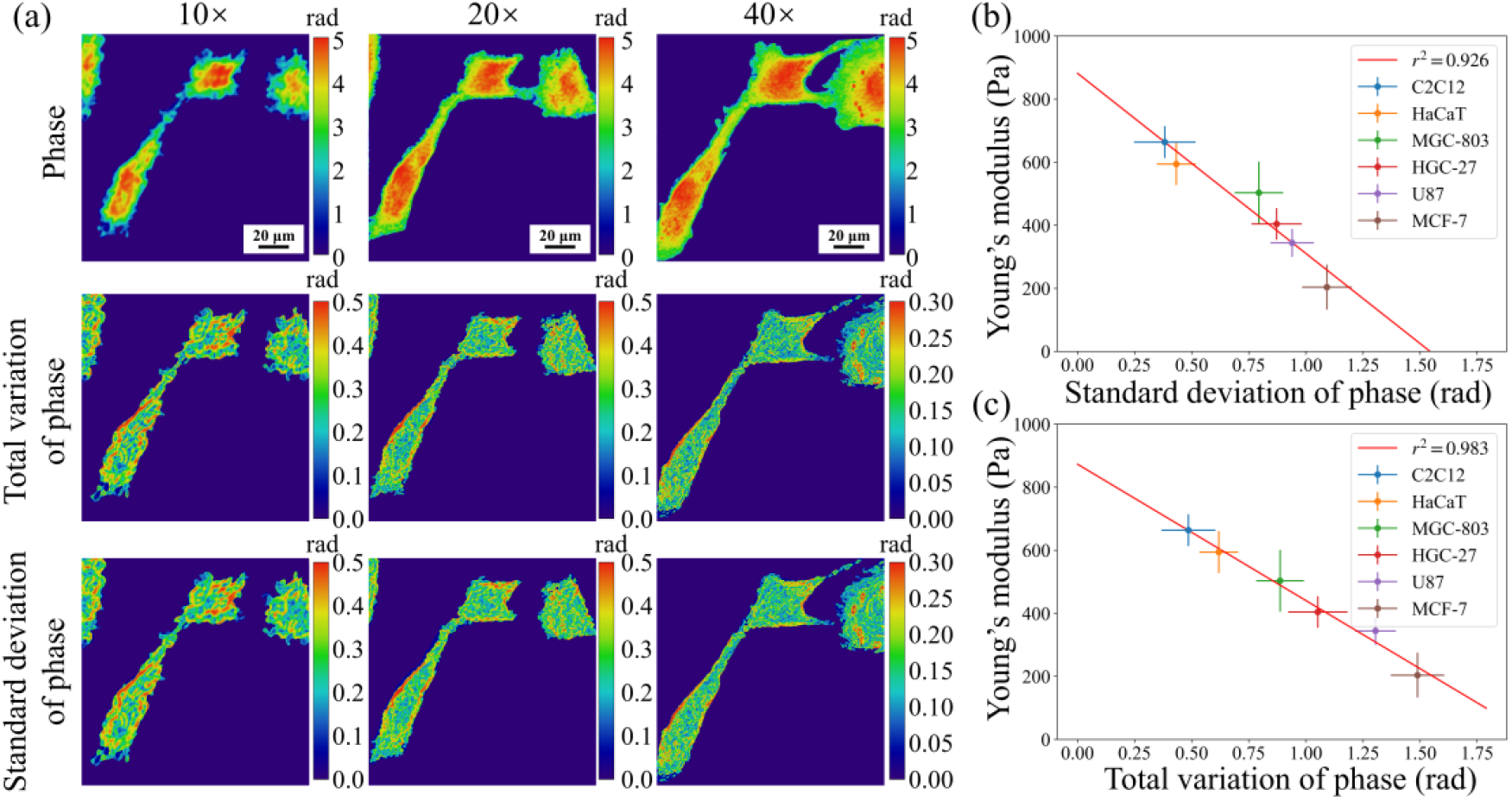
>The performance of both two metrics of phase in high-throughput low-resolution phase images. (a) The comparisons of phase, total variation of phase, and standard deviation of phase of C2C12 cells in different objective lens (OL). (b) The inverse relationship between Young’s modulus and standard deviation of phase in 10× OL. (c) The inverse relationship between Young’s modulus and total variation of phase in 10× OL.

### Mapping the cellular stiffness onto the cell monolayer during collective cell migration using total variation of phase

We used total variation of phase to characterize the spatial distribution of cellular stiffness in a cell monolayer during collective cell migration in *in vitro* wound healing assay. Three types of edges of the wound frontier were imaged using iQPM, which were shown in Figure 5a-c. We further calculated the total variation of phase of three different gap edges and the gradient vectors of Young’s modulus of cells in the monolayer (Figure 5d-f). The total variation of phase of boundary cells was significantly higher than those of inner cells. Additionally, in Figure 5d and Figure 5e, we observed leader cells protruding out of the wound frontiers ^36^, and those leader cells exhibited notably higher total variation of phase (lower Young’s modulus) than other cells. Moreover, in regions where a leader cell is present, we observed that the gradient vectors of Young’s modulus in collective HaCaT cells are distinctly oriented toward the wavy edge formed by the leader cell. To further analyze the relationship between cellular Young’s modulus and its distance from frontier, we draw five 25-μm stripes in Figures 5a-c and calculated the mean total variation of phase for all cells within each stripe. The results were shown in Figure 5g-i. The total variation of phase of collective HaCaT cells was inverse related to the distance from the frontier until it increased to a constant. Thus, the collective HaCaT Young’s modulus has an inverse relationship from the frontier compared to total variation of phase.

**Figure 5.**
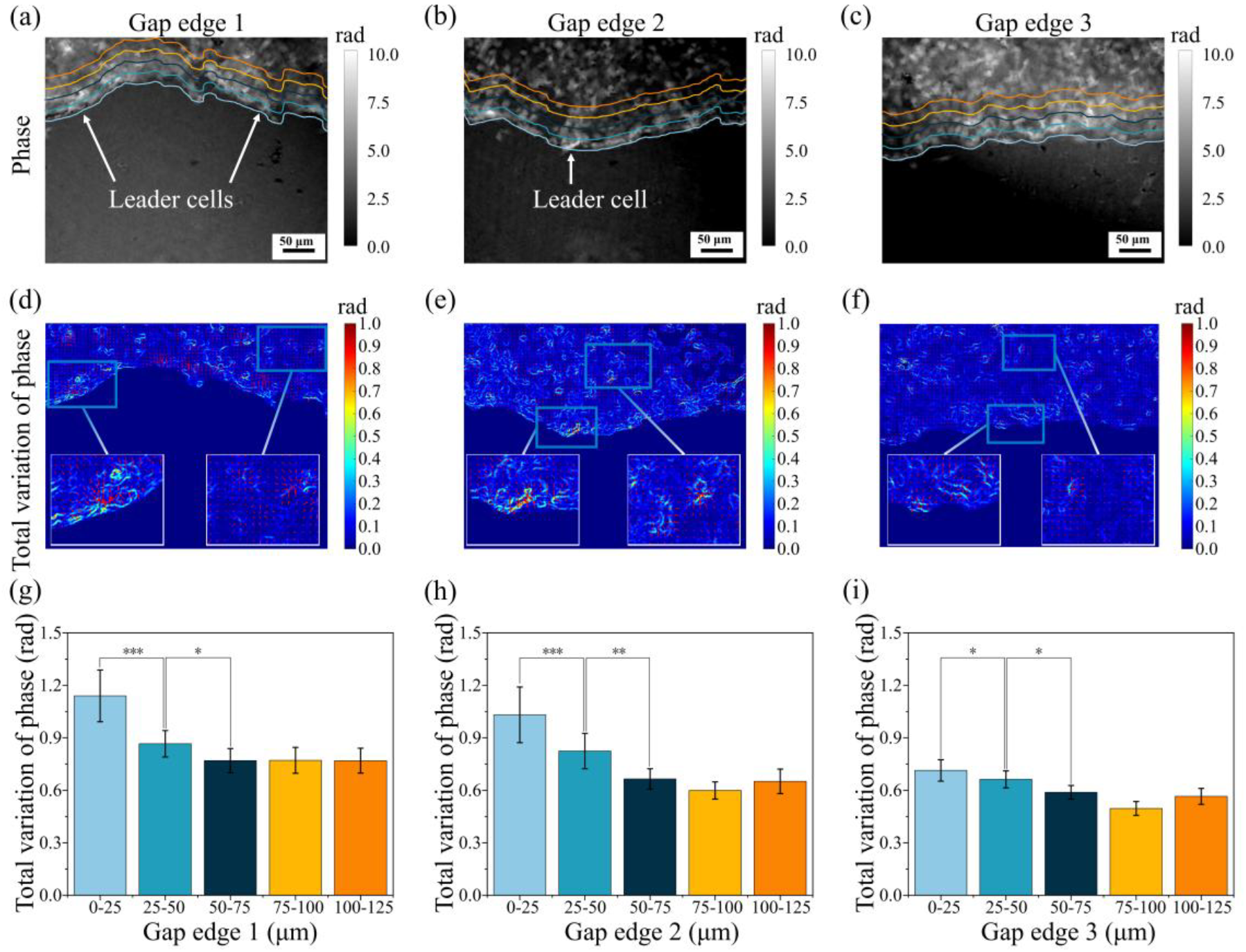
>Using total variation of phase to analyze collective cells migration with 10× OL in about 400μm × 300μm FOV. (a-c) The phase images of collective HaCaT cells migration in different gap edges. (d-f) The total variation of phase images and the gradient vectors of cells of three gap edges. (g-i) The relationship between collective HaCaT cellular Young’s modulus and the distance from the frontier in (a-c), respectively.

To exploit mechanical properties of collective cells migration in single cell resolution, we used iQPM with a 40× OL to scan the collective MCF-7 cells for 6 hours (after scratching and incubating at 37 °C and 5 % CO_2_ for 4 hours) to observe their migration. The quantitative phase images for 6 hours were shown in Figure 6a-c. In high resolution phase images, the leader cell is clear in collective MCF-7 cells. We marked the positions of three different cells (leader cell, follow cell, inner cell) in collective MCF-7 cells in Figure 6a. The calculated total variation of phase images in 6 hours were shown in Figure 6d-f. The total variation of phase of leader cell was significantly increased from Figure 6d to Figure 6f and started to pull following cells for migration. Using the developed tool, we draw gradient vectors of Young’s modulus in collective MCF-7 cells and movements of cellular center-of-mass shifts to reveal relationship between gradient vectors and movements. To eliminate the impact of dead cells on our analysis, we manually excluded data of dead cells. The results were shown in Figure 6g and Figure 6h, the orientations of gradient vectors are similar as its of movement of center-of-mass shifts near the gap edge. However, for some inner cells, the orientations of gradient vectors differ with its of movement. We quantified the changes of total variation of phase from different distance from frontier. To avoid the impact of dead cells, we filtered out the upper 15% and the lower 15% of the total variation of phase and the result was shown in Figure 6i. The near edge cells show larger total variation of phase than others and the difference is increasing as migration. And the changes of total variation of three different cells (leader cell, follow cell, inner cell) was shown in Figure 6j. The leader cell softened as migration, and then the follow cell softened later. However, inner cell showed a steadily total variation and Young’s modulus for 6 hours.

**Figure 6.**
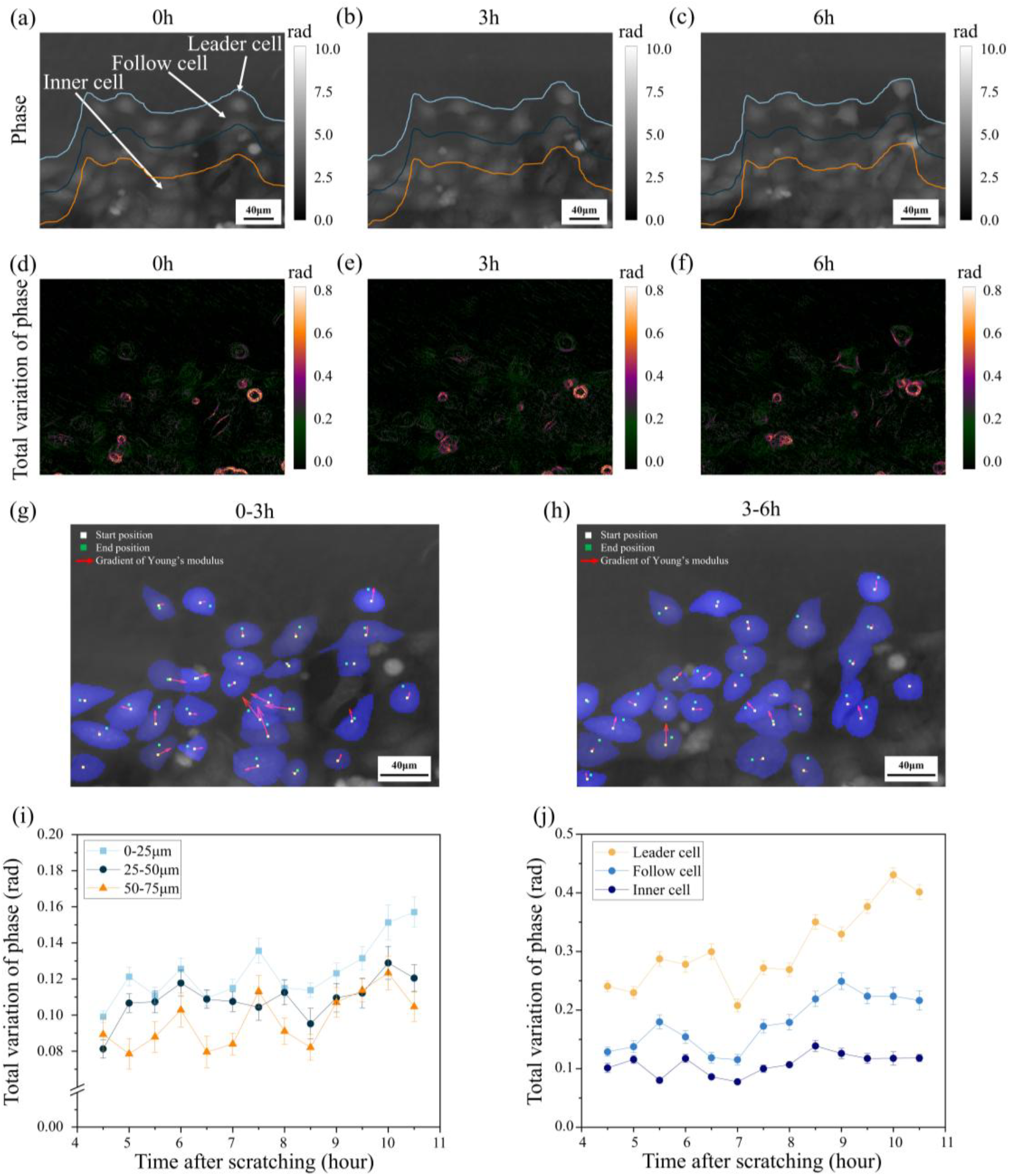
>Using total variation of phase to analyze collective cells migration with 40× OL scanning in about 200μm × 200μm FOV. (a-c) The phase images of collective MCF-7 cells migration during 6 hours. (d-f) The total variation of phase images of collective MCF-7 cells migration during 6 hours. (g) The gradient vectors of Young’s modulus in collective MCF-7 cells show similar directories as cells movements from 0 hour to 3 hour. (h) The gradient vectors of Young’s modulus in collective MCF-7 cells show similar directories as cells movements from 3 hour to 6 hour. (i) The relationship between collective MCF-7 Young’s modulus and the distance from the frontier during 6 hours. (j) The changes of total variation of phase between three different cells during 6 hours.

## DISCUSSION

Integrating AFM and iQPM investigations, we found the total variation of phase linked to the cellular stiffness via the fluctuations of cellular height. The AFM results indicated that the cellular stiffness is inversely related to the fluctuations of cellular height. To overcome the low efficiency of AFM, we imaged the cells with iQPM and found an inverse relationship between the total variation of phase and the cellular Young’s modulus. Integrating this finding with cell segmentation algorithm, we developed a high throughput method to characterize cellular stiffness with single cell resolution. Using the developed method, we further investigated the role of cellular stiffness in collective cell migration. We found that the cells at the wound frontier were much softer than cells in the inner region.

Both standard deviation and total variation of cellular height successfully characterized cellular Young’s modulus qualitatively. AFM have been widely used to characterize the morphology and stiffness of cells. Using peak force tapping mode of AFM, we can get the height and stiffness information at the same time. In the AFM results, we found an inverse relationship between the cellular stiffness and the fluctuation of cellular height. According to the theory of cellular tensegrity, the cytoskeleton structures support the cells body mechanically ^37^. The AFM results in Figure 1 a and b demonstrated the fiber-like structures, which are highly likely F-actin stress fiber. This is validated in AFM results and F-actin fluorescence images of the same cell (Figure 1 g, h, m). Interpreting these results using tensegrity theory, the regions with high density of F-actin stress fiber are smoother and stiffer at the same time. The total variation and standard deviation of height, the quantitative measurement of height fluctuation, demonstrated an inverse relationship with cellular stiffness. These results collectively established the relationship between total variation and standard deviation of cellular height and cellular stiffness.

Though it has nanometer scale resolution, AFM experiments are time consuming and low throughput. We further imaged cells using iQPM for characterizing mechanical properties in fast and high throughput manner. The optical beam experienced phase delay due to the difference of refractive index between the medium and cells. We found the standard deviation and total variation of phase is inversely proportional to the cellular Young’s modulus (Figure 3). If we assumed the *Δn*(*x, y*) in Equation (3), at **Materials and Methods** sections, as a constant in local cellular area, the phase difference between the sample beam and the reference beam reflects the cellular height information. The fluctuations of phase, quantitatively evaluated by their standard deviation and total variation, characterize cellular height, further the cellular stiffness, in a high throughput and fast manner. Like the total variation of cell height case, the total variation of phase has a higher resolution of representing cellular Young’s Modules. Two reasons may account for this. Firstly, standard deviation calculates the fluctuations relative to the mean in the 3×3 pixel^2^ area. It is a low-pass filter for the phase signals, which reduces the differences between signals. Total variation, however, directly calculates the differences between neighboring pixels. Secondly, total variation decomposes the difference into two directions in the 2-D plane, which makes it more directional and better suited for capturing cytoskeletons structures.

One concern of the iQPM method is that we are not sure how deep the cytosol we are characterizing. Is the vicinity of cell membranes or the whole cytosol? To test this, we treated the cells with Y27632, a Rho inhibitor. Compared with control group, the F-actin in the whole cytosol was changed and the total variation of phase identified this stiffness change. This means that iQPM can capture the tiny fluctuations of the cellular height induced by Brownian force from the medium molecules.

Compared with the AFM, the iQPM method has two advantages. First, it is fast and high throughput. Scanning a cell using AFM cost time in minutes, while taking an iQPM images cost less than 1 second. The AFM scans or measures cells features in sequence manner, while all the cells in one iQPM image can be characterized in parallel manner. The throughput can be further increased by using low magnificent objective lens (Figure 4). Second, the AFM cantilever, either with a tip or micro-particle, exerts a large force onto cells. This force deforms the cells and induces cellular reactions, both of which affect the conditions of cells. The iQPM, however, are non-invasive. It affects cellular behaviors in a minimal manner.

The total variation of phase essentially represents the absolute value of its spatial derivative. The relationship between phase (*p*) and Young’s modulus (*s*) in spatial plane (*r*) can be modeled as shown in Equation (1), where *k* and *C* are the constant coefficients.

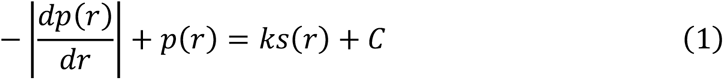

After further operation on Equation (1) (more details are shown in Formula S1 of the Supporting Information), we satisfied an inverse nonlinear relationship between disorder strength of phase and Young’s modulus as *Wax et. al* priorly got ^23,24^, as shown in Equation (2), where *C*_*n*_ is 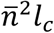 and *C*^″^ is the constant.

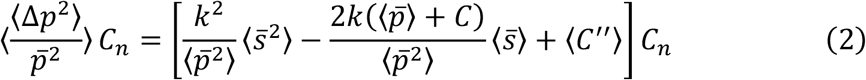

We propose that, under normal conditions, the relationship between disorder strength and Young’s modulus lies within a finite, monotonically decreasing interval where *s*(*r*) > 0. This further confirms that both the spatial distribution of phase in adherent cells and disorder strength of phase characterize the cytoskeleton distribution. This provides a potential framework to further explore the quantitative relationship between phase and cellular mechanical properties.

In recent years, the biological significance of mechanical characters of cells has been widely accepted. Leader-followers coordinated migration group were identified in various collective cell migration scenarios ^38^. The emergent leader cells influence collective cell behaviors through cell-cell interactions and force transmission ^39,40^. These groups lead the cell monolayer migrating forward ^41,42^. Leader cells are usually distinct from other cells by their unique morphologies ^41^. the mechanical character of leader cells, which indicates the molecular structures inside, has been less studied. Will the leader cells be mechanically distinct from other phenotype cells in collective cell migration?

Using the proposed method, we thus characterized the cellular stiffness in collective cell migration in wound healing assay. We found that the wound frontier region of the cell monolayer is much softer than the inner region of the monolayer. In addition, the gradients of collective cellular Young’s modulus point toward the wound edge in Figure 5. The gradients of cellular Young’s modulus near the edge is similar as the cell movement of center-of-mass shifts in Figure 6h-i. This is measuring the cellular stiffness in a large monolayer while the cells are migrating. The softer cells are more invasive ^38^. As the cells migrating, the leader cells and the follower cells are getting softer and softer. This demonstrates the developing process of the actively migrating leader cells and wound frontier region. The directions of cellular stiffness gradients and cellular movements exist discrepancy. The reason might be these cells were not influenced by leader cells. As for single cell in Figure 6h and Figure 6i, without cell-cell interactions, there is no force transmission. Thus, single cell movement is mainly regulated by the mechanical properties of extracellular matrices through mechanotransduction ^43,44^. As for inner cells, the influence of leader cell and force transmission may be constrained by distance, making it difficult to equate movements to gradients of Young’s modulus ^45,46^.

To summarize, the metrics detailed in this study are resolution-efficient and can be directly applied to images with large field of view containing many cells. Combining this approach with the high temporal resolution of iQPM, the two metrics potentially enable high-throughput in situ measurement of cellular Young’s modulus. This technology has broad applications in studying collective cell migration, which is a critical process in many physiological and pathological phenomena. In collective cell migration, some cells at the leading edge transform into leader cells, exhibiting significant polarization characteristics and pulling follower cells to drive migration. However, the mechanisms underlying leader cell formation and the coordination between cells remain unanswered ^47^. Prior studies have described the underlying mechanisms of leader cell formation based on intrinsic properties ^48^. This work contributes to understanding the formation mechanisms of leader cells and collective cells behaviors from the perspective of physical characteristics, particularly the Young’s modulus. It will help elucidate the mechanical signatures of collective invasion for cancer and provide new insights to therapy imperfect wound healing in chronic diseases ^2,49,50^.

## MATERIALS AND METHODS

### Instrument

The migration experiments were conducted using an off-axis quantitative phase microscope (NHQLive Prototype, Shenzhen BJR Biomedical Tech Co. Ltd, China) equipped with an objective lens (10×, NA 0.3, Olympus, Japan), providing a field of view (FOV) of about 500 μm × 500 μm. For other experiments, the NHQLive Prototype was modified by replacing the objective lens (40×, NA 0.65, Olympus, Japan) and utilizing a USB camera (1280×1024, FL3-U3-13Y3M-C, FLIR (Point grey)), resulting in a reduced FOV of about 100 μm × 100 μm. More details were shown in Figure S2.

AFM measurements were performed using the JPK NanoWizard II (JPK Instruments, Germany) mounted on an inverted optical microscope (20×, Axioobserver D1, Nikon, Japan). The sensitivity of the laser detection system was calibrated by measuring the slopes of force-distance curves acquired on a plastic dish, whereas the spring constant of the cantilever was calibrated by the thermal noise method following the instruction provided by JPK Instruments. Aligned to the center of the targeted cell, the AFM tip indented to a depth of 1.0 μm at a rate of 0.3 μm/second. The Young’s modulus was extracted from the force-distance curves using the Hertz model built into the JPK Data Processing software (JPK DP, JPK instrument, Germany). The Young’s modulus of all points within the entire cell area were averaged to obtain the mean Young’s modulus for that cell. For each cell type, we measured approximately 10 cells and calculated their mean Young’s modulus.

### Cell Experiments

Mouse myoblast cells (C2C12), human immortal keratinocytes (*HaCaT)*, gastric cancer cells (HGC-27), glioblastoma cell (U87), human breast cancer cells (MCF-7), and human gastric cancer cells (MGC-803) were cultured in Dulbecco’s Modified Eagle Medium (Gibco, USA) supplemented with 10% fetal bovine serum and 1% penicillin-streptomycin (Gibco, USA). All cells were incubated at 37 °C in a humidified atmosphere with 5% CO_2_. The medium was refreshed daily. Actin-Tracker Green-488 (Beyotime, China) and Hoechst 33342 (Beyotime, China) were used to stain the cells. ECLIPSE Ti2-E (Nikon Instruments Inc., Japan) was used to image the stained C2C12 cells. And in wound healing experiments, 70 ul cell suspension of HaCaT was seeded into the ibidi Culture-Insert 2 Well (ibidi, Germany) on the 60 mm culture dish and incubated at 37 °C and 5 % CO_2_. After reaching 90-100 % confluency, the Culture-Insert 2 Well was gently removed and the cells washed three times with phosphate-buffered saline (PBS).

To evaluate the impact of the cytoskeleton on fluctuations of optical phase delay, we treated C2C12 cells with Y-27632 (SC0326-10mM, Beyotime, China), a Rho-associated protein kinase (ROCK) inhibitor to deliberately decrease cellular stiffness ^33,51^. We prepared a 200 µM stock solution of Y-27632 by dissolving 10 mM × 0.2 ml of the Y27632 solution in 10 ml of dimethyl sulfoxide (DMSO, Sigma Aldrich, Germany). The C2C12 cells were cultured in medium for two days, followed by 24 hours of starvation in 1ml of serum-free DMEM medium. The cells were then treated with 30 µM Y-27632 in cell culture medium for 30 minutes ^51^. After treatment, the Y-27632-treated C2C12 cells were washed twice with PBS before experiments.

### Standard Deviation and Total Variation

The standard deviation of pixels of an image are widely used to represent the signal fluctuations ^34^. Considering the assumption that fluctuations of thickness are primarily generated by the cytoskeleton structure and its directionality, we further introduced total variation to analyze these fluctuations ^52^. According to following equation, the optical phase delay is the product of cellular thickness *d*(*x, y*) and intercellular refractive index *Δn*(*x, y*).

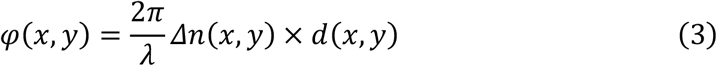

When the objective lens magnification and the resolution are sufficient, the cellular thickness between adjacent pixels in the quantitative phase image, such as (*x, y*) and (*x* + 1, *y*), are approximately same ^23^. However, for high-throughput measurements, the differences of *d*(*x, y*) between adjacent pixels cannot be ignored. These thickness differences (0-10 μm), (add unit)which is significantly larger than the differences of refractive index across different cellular parts (normally on the order of 10^−2^) ^53^. We assumed that *Δn*(*x, y*) within the cell at low resolution was constant, as the variations of dry mass within a cell were sufficiently small.

We directly calculated these two metrics of the phase in each 3×3 pixel^2^ area or in two orientations. Despite both standard deviation and total variation representing fluctuations, there are still differences between them. Standard deviation is a statistical matrix. As shown in following equation, *p*(*x, y*) is the optical phase delay at 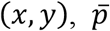 is the mean value of the optical phase delay in the 3×3 pixel^2^ area, and *n* is 9 in 3×3 pixel^2^ area.

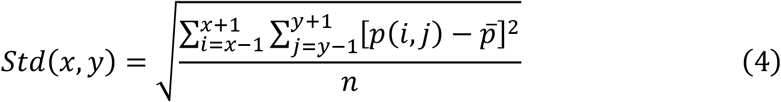

In contrast, total variation is the absolute value of the spatial derivative, as shown in following equation.

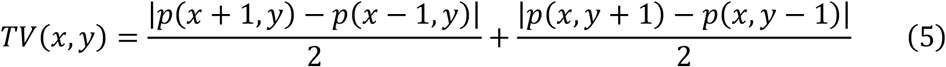

### Statistical Analyses

Statistical analyses were conducted in Python 3.10, supplemented by various Python libraries, including Pandas 2.0.2, Matplotlib 3.7.1, Numpy 1.24.3, and Scipy 1.10.1. For each cell type, the mean of the measured values for 10 cells was calculated. These data were linearly fitted, and the r-squared values were calculated. Statistical significance was evaluated using a two-tailed student’s t-test, and the p-value was provided in the results.

For gradient of collective cellular Young’s modulus, we treated each 16×16 pixel^2^ as a unit and calculated the mean total variation of phase of each unit. We calculated the gradient of total variation between another unit (within a specified distance for about 150μm range from the target unit) and the target unit (considering the impact of distance, the calculated gradient is divided to the square of distance between two cell). Add all gradients for the gradient of total variation of one point. The gradient of Young’s modulus is the reversing gradient of total variation of phase. By repeating above process, we draw the gradient map of collective cellular Young’s modulus. For cell resolution gradients analysis, we used ASF-YOLO to segment cells in phase images (Figure S1), and then calculated start points and end points of center of mass to demonstrate movements. We then calculated the gradient between another cell (within a specified distance of center of mass for about 150μm range from the target cell) and the target cell (considering the impact of distance, the calculated gradient is divided to the square of distance between two cell). Add all gradients for the gradient of total variation of one cell. We then added a bias (gradient of total variation between a cell and the extracellular matrix, which is 10^−6^ rad) to the calculated gradient. The gradient of Young’s modulus is the reversing gradient of total variation of phase.

## Supporting information

The supplemental figure and prove

## Data availability

The authors declare that the data supporting the findings of this study are available within the paper and its supplementary information files.

## AUTHOR CONTRIBUTIONS

Min Long and Yongliang Yang conceived the idea, designed the experiments, and edited the manuscript. Zhenghua Wang did all the experiments and drafted the manuscript. Zebin Wang, Deyu Sun, and Yujie Nie helped the experiments related with QPI. Jialin Shi, Renjie Zhou, and Lianqing Liu helped designed the experiments, discussed the results, and edited the manuscript.

## CONFLICT OF INTEREST STATEMENT

All the authors have no conflict of interest.

## Notes

### Competing Interest Statement

The authors have declared no competing interest.

### Funding

- National Natural Science Foundation of China: 62273338

