## Supplementary material for "Total variation of quantitative phase map reveals cellular Young’s modulus": The supplemental figure and prove

Yongliang Yang: 114 Nanta Street, Shenhe District, Shenyang City, Liaoning, China

 , +86-24-23974593

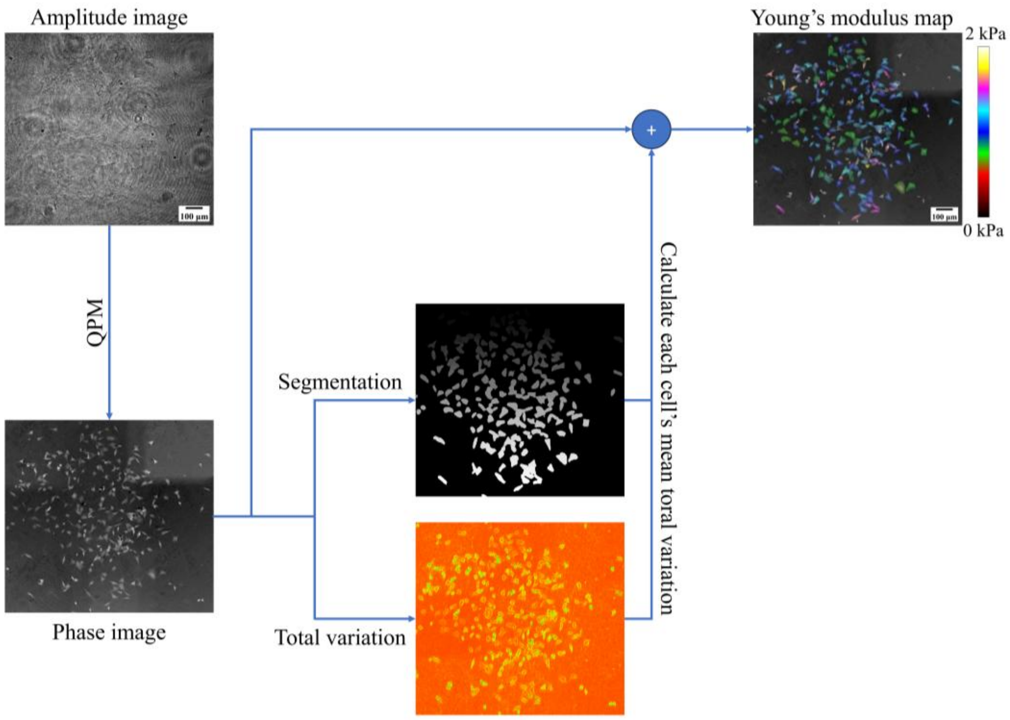


**Figure. S1**. Diagram of the proposed automated high throughput cellular Young's modulus measuring tool. We employed quantitative phase microscopy to image quantitative optical phase delay of cells. Then by combining a deep learning model and the phase image, the spatial location of each cell was captured. By integrating the cell location with the proposed total variation of phase, the map of mean total variation of phase was calculated to characterize the Young's modulus of collective cells. For well visualization, the map of cellular Young's modulus was combined with the phase image to yield a clear result.

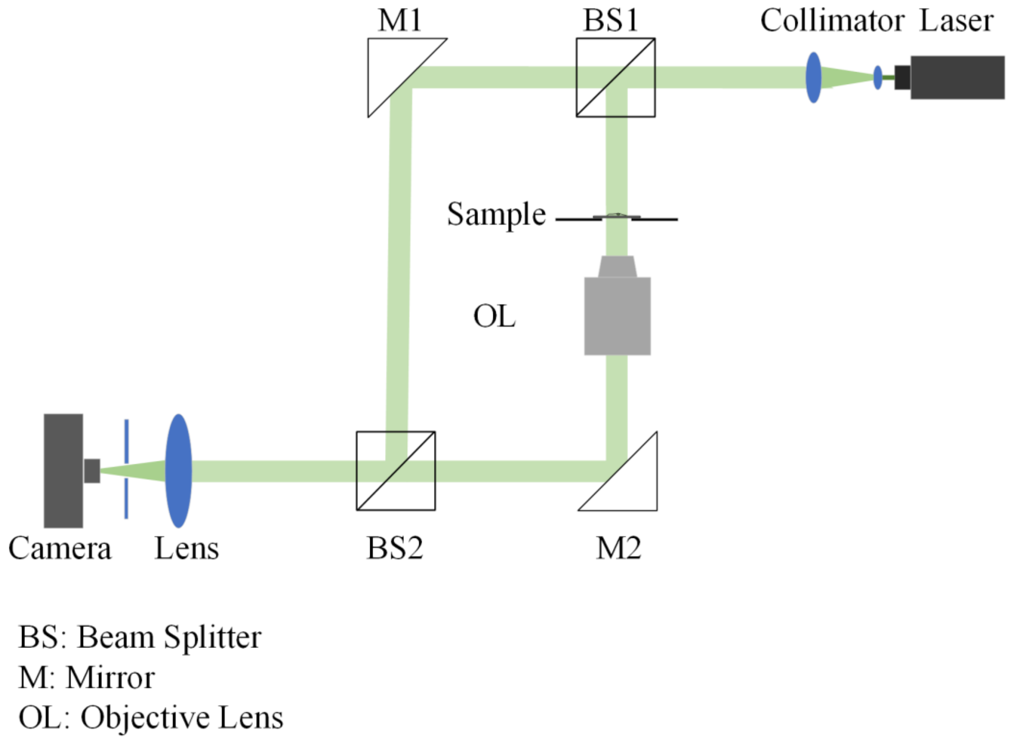


**Figure. S2**. Experiment setup of iQPM.

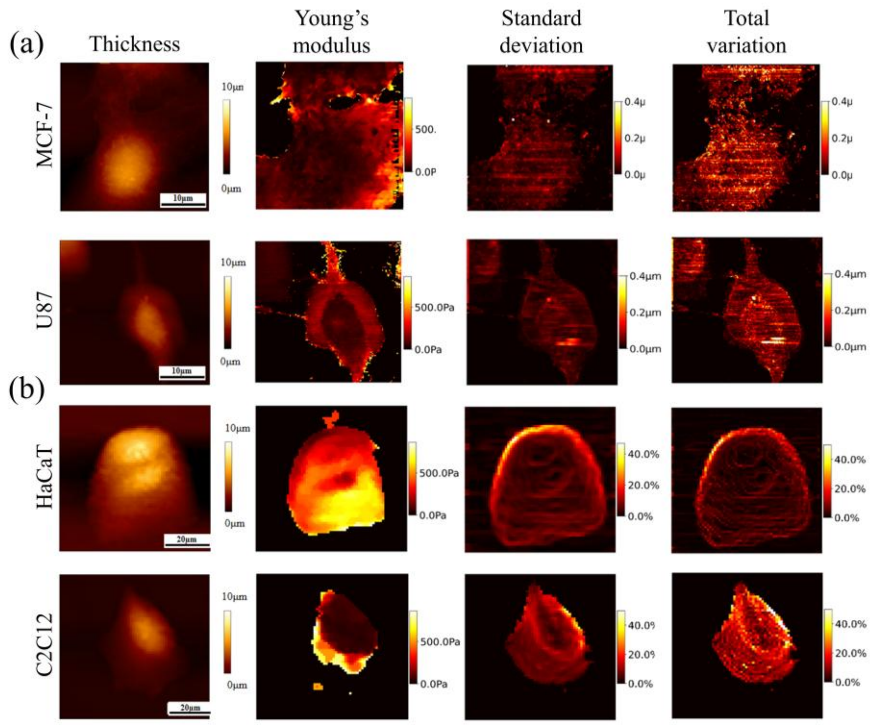


**Figure. S3**. The thickness images, Young's modulus images, calculated standard deviation of thickness images, and calculated total variation of thickness images. (a) The thickness images, Young's modulus images, calculated standard deviation of thickness images, and calculated total variation of thickness images in whole cell. (b)

The thickness images, Young's modulus images, calculated percentages of standard deviation of thickness images, and calculated percentages of total variation of thickness images in whole cell.

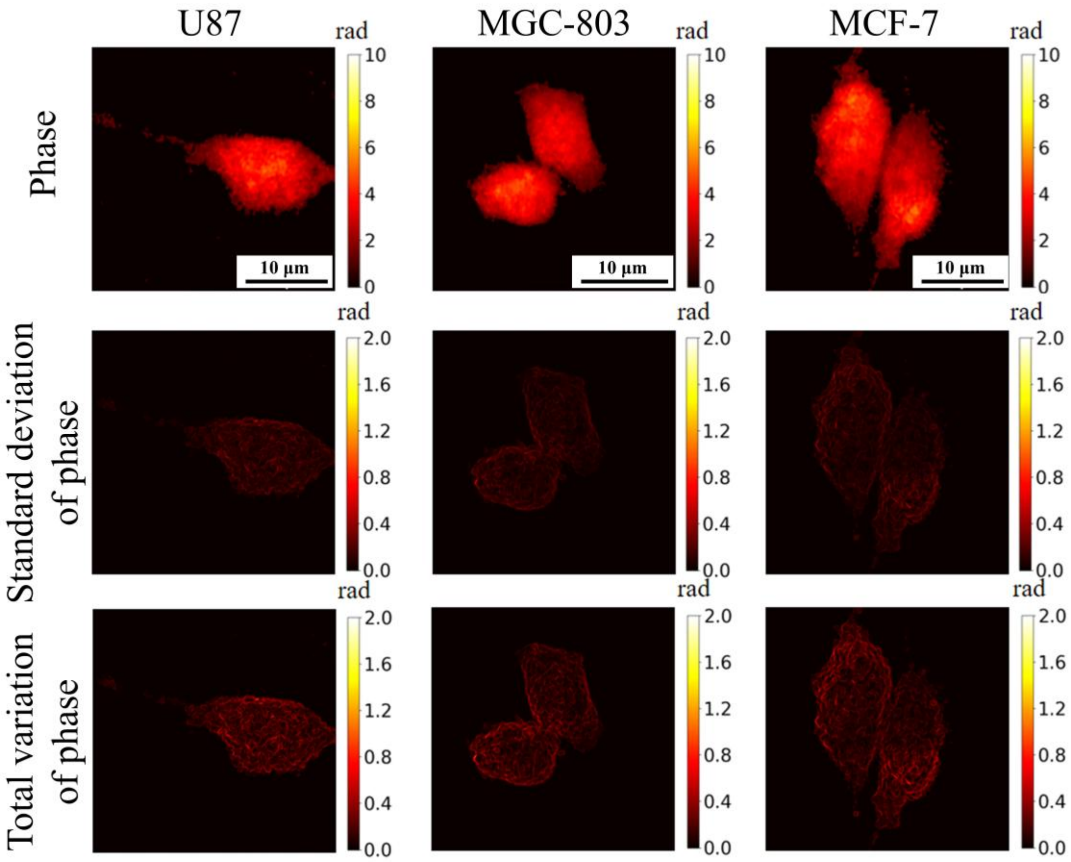


**Figure. S4**. The phase images, standard deviation of phase images, and total variation of phase images of more types of cells.

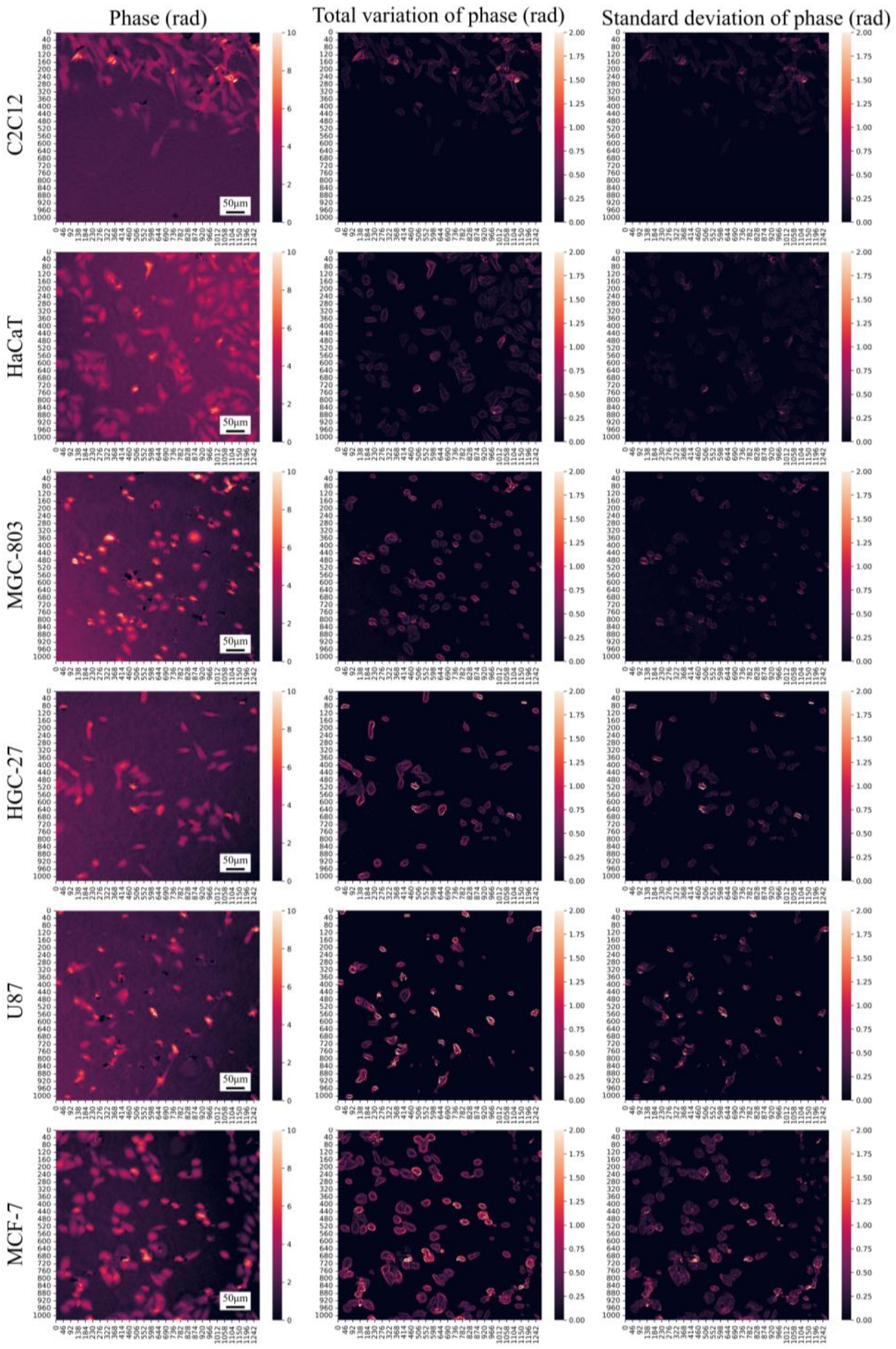


**Figure. S5**. The phase images, standard deviation of phase images, and total variation of phase images of all six types of cells in 10× OL.

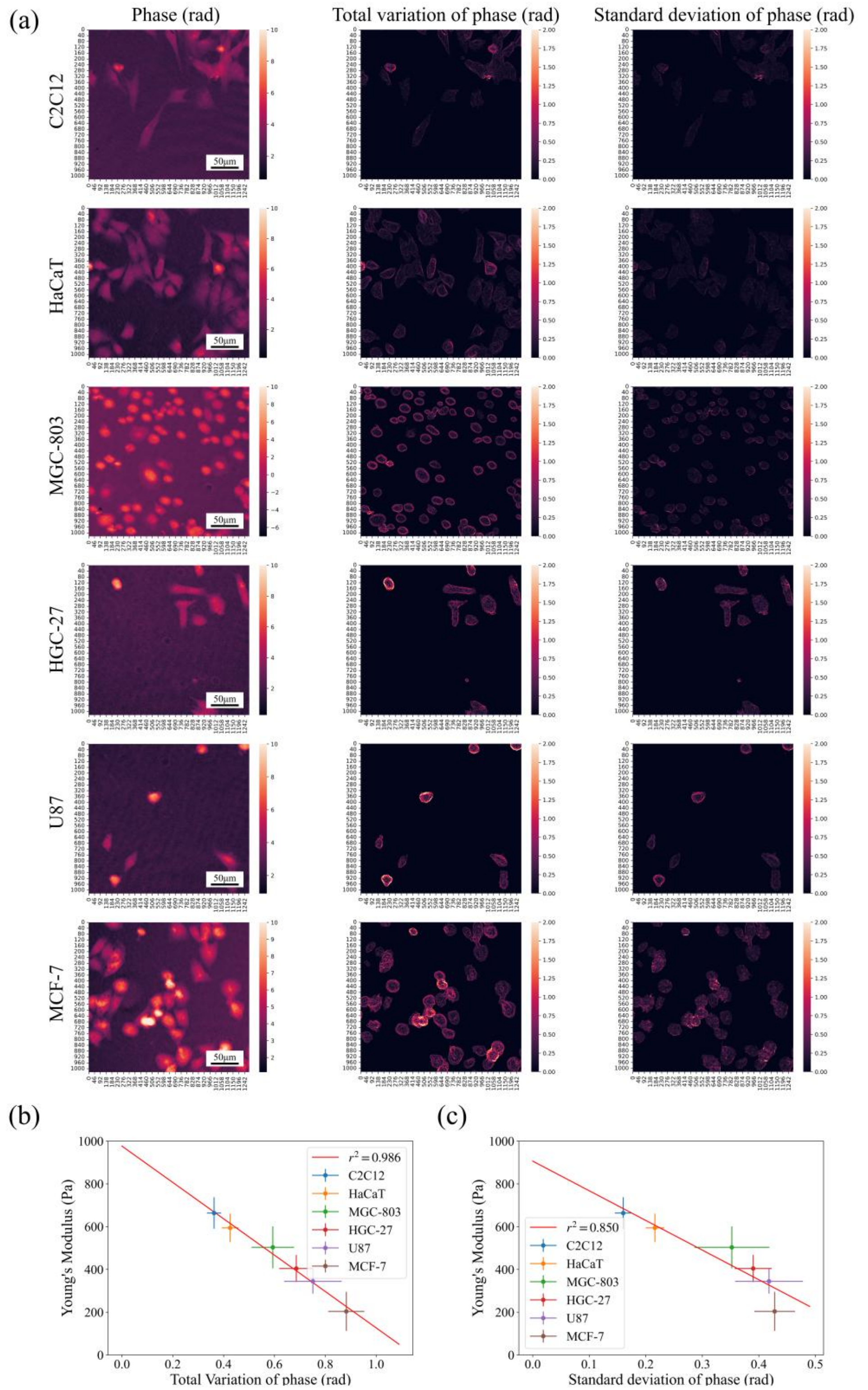


**Figure. S6**. The phase images, standard deviation of phase images, and total variation of phase images of all six types of cells and their inverse relationship images in 20× OL. (a) The phase images, standard deviation of phase images, and total variation of phase images of all six types of cells. (b) The inverse relationship between Young's modulus and total variation of phase. (c) The inverse relationship between Young's modulus and standard deviation of phase.

**Formula S1. Proof**

$$-\left|\frac{dp(r)}{dr}\right|+p(r)=ks(r)+C \qquad (4)$$

If we ignored $|\cdot|$, the above equation was written as:

$$-\frac{dp(r)}{dr}+p(r)=ks(r)+C \qquad (S1)$$

$$-dp(r)+p(r)dr=ks(r)dr+Cdr \qquad (S2)$$

When we integrate Equation S2 over a discrete finite region, $C$ is a constant.

$$-\int dp(r)+\int_{s}^{s+n} p(r)dr=k\int_{s}^{s+n} s(r)dr+nC \qquad (S3)$$

$$-\int dp(r)+n\bar{p}=nk\bar{s}+nC \qquad (S4)$$

We interpret the integration of the derivative as the subtraction between two corresponding sets $P_1=\{p(r)|r_{s+n}>r>r_s\}$ and $P_2=\{p(r)|r_{s+n-1}>r>r_{s-1}\}$. And when region is very small, the $\sum_{r\in P_2} p(r)\approx n\overline{p(r)}$, both $C'$ and $C''$ mean constants.

$$-\sum_{r\in P_1} p(r)+\sum_{r\in P_2} p(r)+n\bar{p}=nk\bar{s}+nC \qquad (S5)$$

$$\sum_{r=r_s}^{r_{s+n}}(p(r)-\bar{p})=-nk\bar{s}+n\bar{p}+nC \qquad (S6)$$

$$\sum_{r=r_s}^{r_{s+n}}\Delta p(r)=-nk\bar{s}+n\bar{p}+nC \qquad (S7)$$

$$\left(\sum_{r=r_s}^{r_{s+n}}\Delta p(r)\right)^2=n^2(k^2\bar{s}^2-2k\bar{s}\bar{p}-2k\bar{s}C+C^2+2C\bar{p}+\bar{p}^2) \qquad (S8)$$

$$n\sum_{r=r_s}^{r_{s+n}}\Delta p^2(r)=n^2k^2\bar{s}^2-2n^2k(\bar{p}+C)\bar{s}+n^2C' \qquad (S9)$$

$$\langle\Delta p^2\rangle=k^2\bar{s}^2-2k(\bar{p}+C)\bar{s}+C' \qquad (S10)$$

$$\frac{\langle \Delta p^2 \rangle}{\bar{p}^2} = \frac{k^2}{\bar{p}^2}\bar{s}^2 - \frac{2k(\bar{p} + C)}{\bar{p}^2}\bar{s} + C'' \tag{S11}$$

A 3×3 region was selected to calculate $\frac{\langle \Delta \mathrm{p}^2 \rangle}{\bar{\mathrm{p}}^2}$.[36, 37] The values of $\frac{\langle \Delta \mathrm{p}^2 \rangle}{\bar{\mathrm{p}}^2}$ over the entire cell were then summed and averaged to $\langle \frac{\langle \Delta \mathrm{p}^2 \rangle}{\bar{\mathrm{p}}^2} \rangle$.

$$\langle \frac{\langle \Delta p^2 \rangle}{\bar{p}^2} \rangle \bar{n}^2 l_c = \left[ \frac{k^2}{\langle \bar{p}^2 \rangle} \langle \bar{s}^2 \rangle - \frac{2k(\langle \bar{p} \rangle + C)}{\langle \bar{p}^2 \rangle} \langle \bar{s} \rangle + \langle C'' \rangle \right] \bar{n}^2 l_c \tag{S12}$$
